# Inhibitory effects of β-caryophyllene on *Helicobacter pylori* infection *in vitro* and *in vivo*

**DOI:** 10.1101/846790

**Authors:** Hyun Jun Woo, Ji Yeong Yang, Min Ho Lee, Hyun Woo Kim, Hye Jin Kwon, Min Park, Sung-kyu Kim, So-Young Park, Sa-Hyun Kim, Jong-Bae Kim

## Abstract

The human specific bacterial pathogen *Helicobacter pylori* (*H. pylori*) is a Gram-negative microaerophilic bacterium and associated with severe gastric diseases such as peptic ulceration and gastric cancer. Recently, the increasing resistance and the emergence of adverse effects make the usage of antibiotics less effectively. Therefore, development of new antimicrobial agent is required to control *H. pylori* infection. In the current study, it has been demonstrated the inhibitory effect of β-caryophyllene on *H. pylori* growth and the protective effect against *H. pylori* infection as well as antibacterial therapeutic effect. β-caryophyllene inhibited *H. pylori* growth via down-regulation of *dna*E, *dna*N, *hol*B and *gyr*A and also down-regulated virulence factors such as CagA, VacA and SecA proteins. β-caryophyllene inhibited expression of several type IV secretion system (T4SS) components including *vir*B2, *vir*B4 and *vir*B8, so that CagA translocation into *H. pylori*-infected AGS cells was decreased by β-caryophyllene treatment. β-caryophyllene also inhibited VacA toxin entry through down-regulation of type IV secretion system (T5SS). *In vivo* experiments using Mongolian gerbils demonstrated antibacterial therapeutic effects of β-caryophyllene. After β-caryophyllene administration, immunohistochemistry (IHC) stain using anti-*H. pylori* antibody showed the antibacterial effect and H&E stain showed the therapeutic effect in treated groups. Hematological data which conformed with histological data support the therapeutic effect of β-caryophyllene administration. Such a positive effect of β-caryophyllene on *H. pylori* infection potently substantiate that this natural compound could be used as a new antimicrobial agent or functional health food to help the patients whom suffering from gastroduodenal diseases due to *H. pylori* infection.

**Author summary:** The inhibitory effect on β-caryophyllene on *H. pylori* growth and the protective effect against *H. pylori* infection as well as antibacterial therapeutic effect have been elucidated in this study. β-caryophyllene inhibited *H. pylori* growth via downregulation of replication machinery of *H. pylori*. β-caryophyllene also downregulated virulence factors such as CagA, VacA and SecA proteins which are necessary for successful colonization and pathogenesis of *H. pylori*. Besides, β-caryophyllene significantly reduced *H. pylori*-induced actin-cytoskeletal rearrangement, vacuolation and apoptosis in AGS cells. In *in vivo* infection model, β-caryophyllene showed splendid therapeutic effect against *H. pylori* infection. In particular, this is the first report that evaluates the toxicological effects of β-caryophyllene administration on Mongolian gerbils. Such a positive effect of β-caryophyllene on *H. pylori* infection potently substantiate that this natural compound could be used as a new antimicrobial agent or functional health food to help the patients whom suffering from gastroduodenal diseases due to *H. pylori* infection.

## Introduction

*Helicobacter pylori* (*H. pylori*) is a Gram-negative, spiral-shaped, microaerophilic bacterium that selectively colonizes in human gastric mucosa [1, 2]. By utilizing virulence factors, *H. pylori* can change the environment and lower the acidity of the stomach, where it impairs the gastric mucosa and alters the gastric acid secretion, thereby disturbing gastric physiology [3]. Compared with uninfected individuals, *H. pylori* infected individuals have 2-8 fold increased risk of developing gastric cancer [4–6]. For these reasons, *H. pylori* has been classified as class I carcinogen by the World Health Organization (WHO) [7]. *H. pylori*-mediated gastric diseases are mostly due to the effect of its virulence factors. Thus, understanding the biological characteristics and mechanisms of their virulence factors might offer a more comprehensive understanding into the pathogenesis of *H. pylori* infection.

Cytotoxin-associated protein A (CagA) is a highly immunogenic protein that is encoded by the *cag* pathogenicity island (PAI). The *cag*PAI genes also encode components of a type IV secretion system (T4SS), which injects CagA protein into host cells [8]. The T4SS apparatus commonly consists of 11 VirB proteins, encoded by the *vir*B1-B11 genes and VirD4 protein [9, 10]. Once translocation, CagA protein is localizes to the inner surface of the plasma membrane via interactions with phosphatidylserine and sequentially phosphorylated at the Glu-Pro-Ile-Tyr-Ala (EPIYA) sequence repeats by the Src and Abl tyrosine kinases [11–13]. The phosphorylated CagA also dysregulates the homeostatic signal transduction of gastric epithelial cells leading the loss of cell polarity, apoptosis and proliferation which involved in chronic inflammation and gastric cancer [14–16].

Vacuolating cytotoxin A (VacA) protein which is known as a pore-forming secreted toxin, found in almost all *H. pylori* strains [17]. VacA secretion is regulated by the type Va secretion system (T5aSS) and depends on the Sec machinery for transport through the inner membrane [18]. Secretion system subunit protein A (SecA) is an intracellular ATPase that provides energy necessary for translocation of *H. pylori* proteins out of the bacterial plasma membrane [18–20]. Following translocation into the host cells, VacA leads the multiple cellular alterations, including vacuolation and cell death [21]. Cellular vacuolation during *H. pylori* infection is considered to disrupt protein trafficking pathways and affect a number of cellular functions, such as intracellular degradation of epidermal growth factor and inhibition of antigen presentation in immune cells [22–24].

DNA replication is the biological process of copying the DNA in the all living organisms. Initiation of DNA replication occurs after binding of initiator protein DnaA to the AT-rich regions on *ori*C. DnaA then forms a complex with other nucleoproteins (DnaBC) and ATPs to make the pre-Replicative Complex [25]. DnaB is DNA helicase, progressively unwinds the double stranded DNA in the 5’-3’ direction [26, 27]. *H. pylori* possesses six DNA polymerse III holoenzyme genes. These include two genes for replicase (DnaE and DnaQ), one for clamp (DnaN) and three for clamp loader (DnaX, HolA and HolB) [25]. The termination of DNA replication takes place when the helicase DnaB on the leading strands reaches a Tus protein [26].

For eradication *H. pylori*, two or more antibiotics are usually prescribed together with a proton pump inhibitor (PPI) and/or bismuth containing compounds [28]. The triple therapy is the most commonly used first-line regimen for eradication of *H. pylori* and consists of amoxicillin, clarithromycin and PPI [28]. However, treatment of *H. pylori* infection is becoming less effective as a result of increasing antibiotic resistance worldwide [29–31]. In particular, clarithromycin resistance in Asia in 2014 was 32.46% according to a recent review by Ghotaslou *et al* [29]. These suggesting that development of a new antibiotic agent which alternatively targeted approach to eradicate *H. pylori* would be needed [32].

β-caryophyllene is a volatile bicyclic sesquiterpene compound which exists as a mixture of mainly β-caryophyllene and small amounts of α-humulene [33]. It is easily found in the essential oils of many edible plants such as cloves (*Eugenia caryophyllata*) [34], oregano (*Origanum vulgare*) and cinnamon (*Cinnamomum* spp.) [33, 35, 36]. Recent approval of β-caryophyllene as the food additives and flavoring agent by the Food and Drug Administration in USA (FDA) and the European Food Safety Authority (EFSA) generated the interests among scientific community to explore its additional therapeutic benefits [37, 38]. Numerous reports showed inhibitory effects of β-caryophyllene against bacteria, virus and fungi [39–42].

Recently, natural compounds derived from medicinal plants seem to be an important source of antimicrobial agents. Because of the increasing resistance and the emergence of adverse effects, the usage of antibiotics and antibacterial therapeutics is becoming less effective. Therefore, inhibitory effect of β-caryophyllene on *H. pylori* growth and its inhibitory mechanisms were investigated in this study. Furthermore, it is aimed to discover a new antimicrobial agent to eradicate *H. pylori,* or identify a new functional health food which can reduce the virulence of *H. pylori* in infected gastric cells *in vitro* and *in vivo*.

## Results

### 1. Inhibitory effect of β-caryophyllene on the growth and expression of replication genes of *H. pylori*

As the inhibitory effect of β-caryophyllene on *H. pylori* was priorly evaluated by screening test using the disc diffusion assay. Clear inhibitory zones were observed around 10, 50 and 100 μg discs and the diameters of the inhibitory zones were 17, 21 and 23 mm, respectively (Fig 1A). As the inhibitory effect was confirmed, the minimal inhibitory concentration (MIC) of β-caryophyllene against *H. pylori* was determined by the broth dilution test. The MIC of β-caryophyllene against *H. pylori* was determined as 1,000 μg/mL (Fig 1B). These results suggest that β-caryophyllene has an inhibitory effect on *H. pylori* growth.

**Fig 1.**
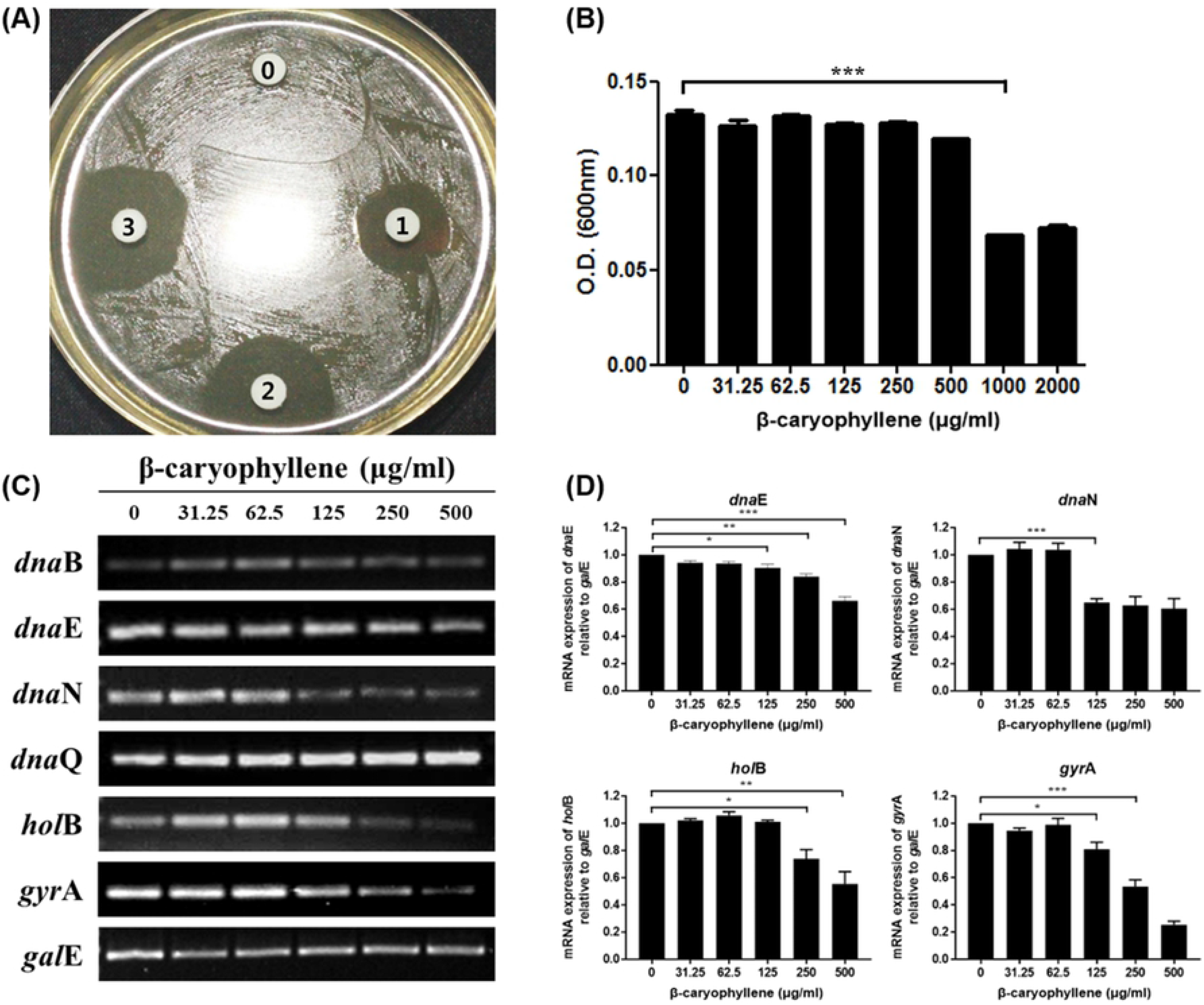
Anti-bacterial activity of β-caryophyllene against *H. pylori* and downregulation of replication-related genes of *H. pylori*. (A) Growth inhibitory activity of β-caryophyllene against *H. pylori* was confirmed by disc diffusion test. Disk 0, control; disk 1, β-caryophyllene 10 μg; disk 2, β-caryophyllene 50 μg; disk 3, β-caryophyllene 100 μg. (B) Minimal inhibitory concentration of β-caryophyllene against *H. pylori* was determined by broth dilution method. Results from triplicate experiments were analyzed by Student’s *t*-test (\*\**P* < 0.001). (C) The mRNA level expression levels of DNA replication machineries. Constitutively expressed *gal*E was used an internal control. (D) Density of the bands were illustrated as a graph, and the results from triplicate experiments were analyzed by Student’s *t*-test (\**P* < 0.05, \*\**P* < 0.01 and \*\*\**P* < 0.001).

To elucidate how β-caryophyllene inhibits growth of *H. pylori*, expressions of replication genes of *H. pylori* were evaluated. RT-PCR was conducted to confirm the downregulation of the replication genes in β-caryophyllene-treated *H. pylori*, targeting various genes associated with replication (*dna*B, *dna*E, *dna*N, *dna*Q *hol*B and *gyr*A) of *H. pylori*. The result confirmed the down-regulation of *dna*E, *dna*N, *hol*B and *gyr*A mRNA levels in *H. pylori* treated with β-caryophyllene (Fig 1C and 1D). Therefore, these results indicate that inhibitory mechanism of β-caryophyllene against *H. pylori* growth is in part associated with interruption of bacterial replication via down-regulation of *dna*B, *dna*N, *hol*B and *gyr*A genes.

### 2. Suppression of *H. pylori*-induced apoptosis in gastric epithelial cells by β-caryophyllene

Infection of *H. pylori* results in deleterious effects on gastric epithelial cells such as disruption of intercellular junction, cytoskeletal rearrangement, vacuolation and induction of apoptosis. Thus it was evaluated whether β-caryophyllene can alleviate the deleterious effects of *H. pylori* infection on gastric epithelial cells. When β-caryophyllene was treated to AGS cells without *H. pylori* infection, the dose higher than 1,000 μg/mL could reduce cell viability (Fig 2A). Thus, the dose of β-caryophyllene upto 500 μg/mL was used in the following experiments to avoid cytotoxic effect of β-caryophyllene.

**Fig 2.**
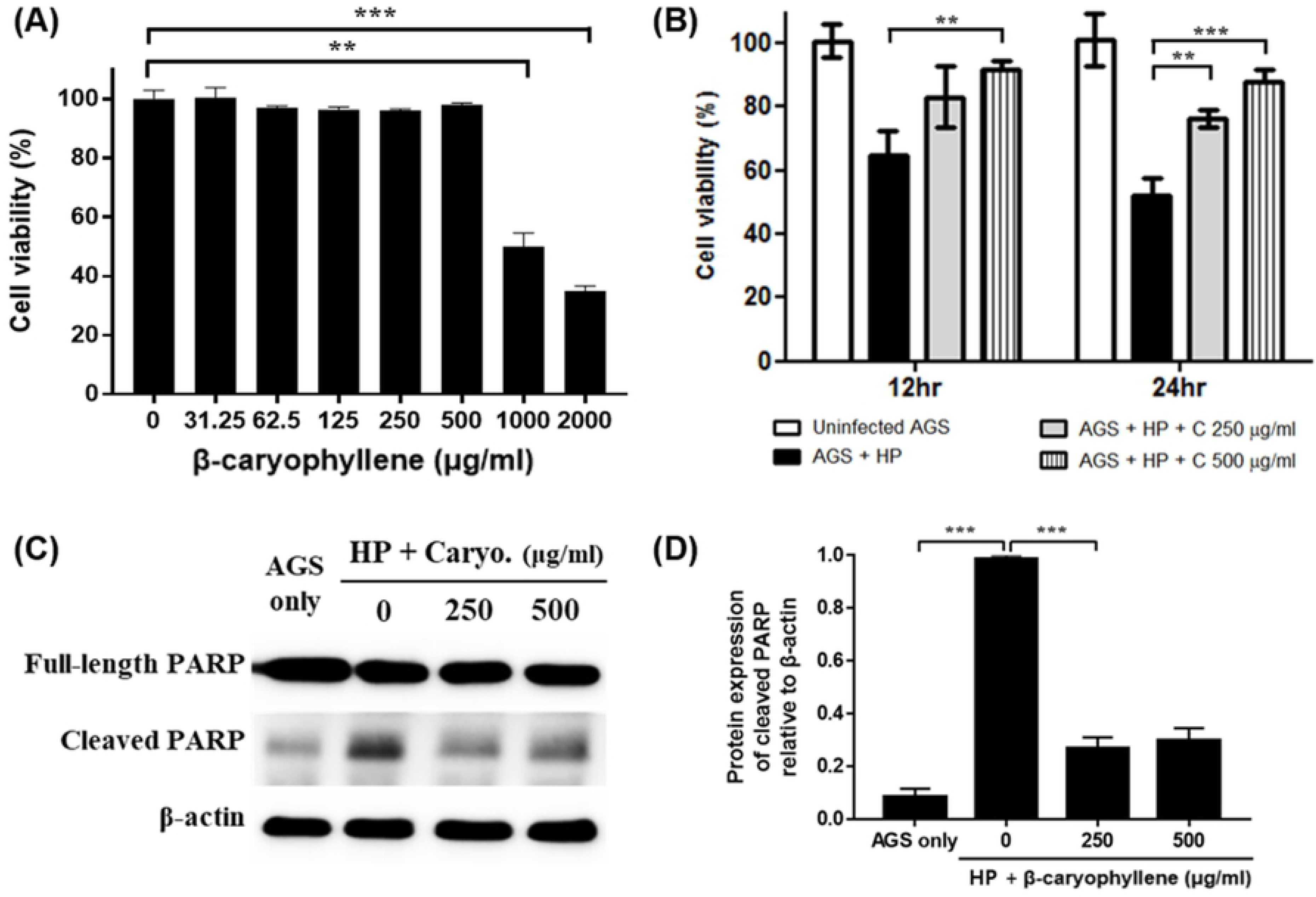
Inhibitory effect of β-caryophyllene on *H. pylori*-infected AGS cell death. (A) AGS cells were treated with the indicated concentrations of β-caryophyllene for 24 h and cell viability was measured by the WST assay. Cell viability of AGS cells was decreased with 1,000 μg/mL or higher dose of β-caryophyllene treatment. AGS cells were infected with *H. pylori* (200 MOI) and treated with β-caryophyllene. (B) After 12 or 24 h, cell viability was measured the WST assay. (C) The cell lysates were assessed by Western blotting to detect a full-length of PARP (116 kDa) and cleaved PARP (89 kDa). β-actin was used as an internal control. (D) Density of the bands were illustrated as a graph. Data in the bar graphs are presented as the mean ± standard error of mean. Data were from the three independent experiments and analyzed by unpaired Student’s *t*-test (\*\**P* < 0.01 and \*\*\**P* < 0.001).

It was also investigated whether β-caryophyllene can alleviate apoptotic cell death of gastric epithelial cells induced by *H. pylori* infection. AGS cells were infected with 200 MOI of *H. pylori* and treated with β-caryophyllene for 12 h or 24 h, and then cell viability was measured by WST assay. *H. pylori* infection reduced cell viability of AGS cells to 51.8% in 24 h, but the reduction of cell viability was alleviated upto 87.6% by 500 μg/mL β-caryophyllene treatment (Fig 2B). In the Western blot, poly ADP-ribose polymerase (PARP) was cleaved in AGS cells by *H. pylori* infection indicating induction of apoptosis. The *H. pylori*-induced PARP cleavage was inhibited by β-caryophyllene treatment (Fig 2C and 2D). Furthermore, annexin V-FITC stain was performed to confirm the presence of apoptosis in the same condition. The result analyzed by flowcytometry also showed that apoptosis induced by *H. pylori* infection was alleviated in AGS cells by β-caryophyllene treatment (Fig 3A and 3B). These results collectively suggest that β-caryophyllene inhibited apoptosis and alleviated cell death in AGS cells infected with *H. pylori*.

**Fig 3.**
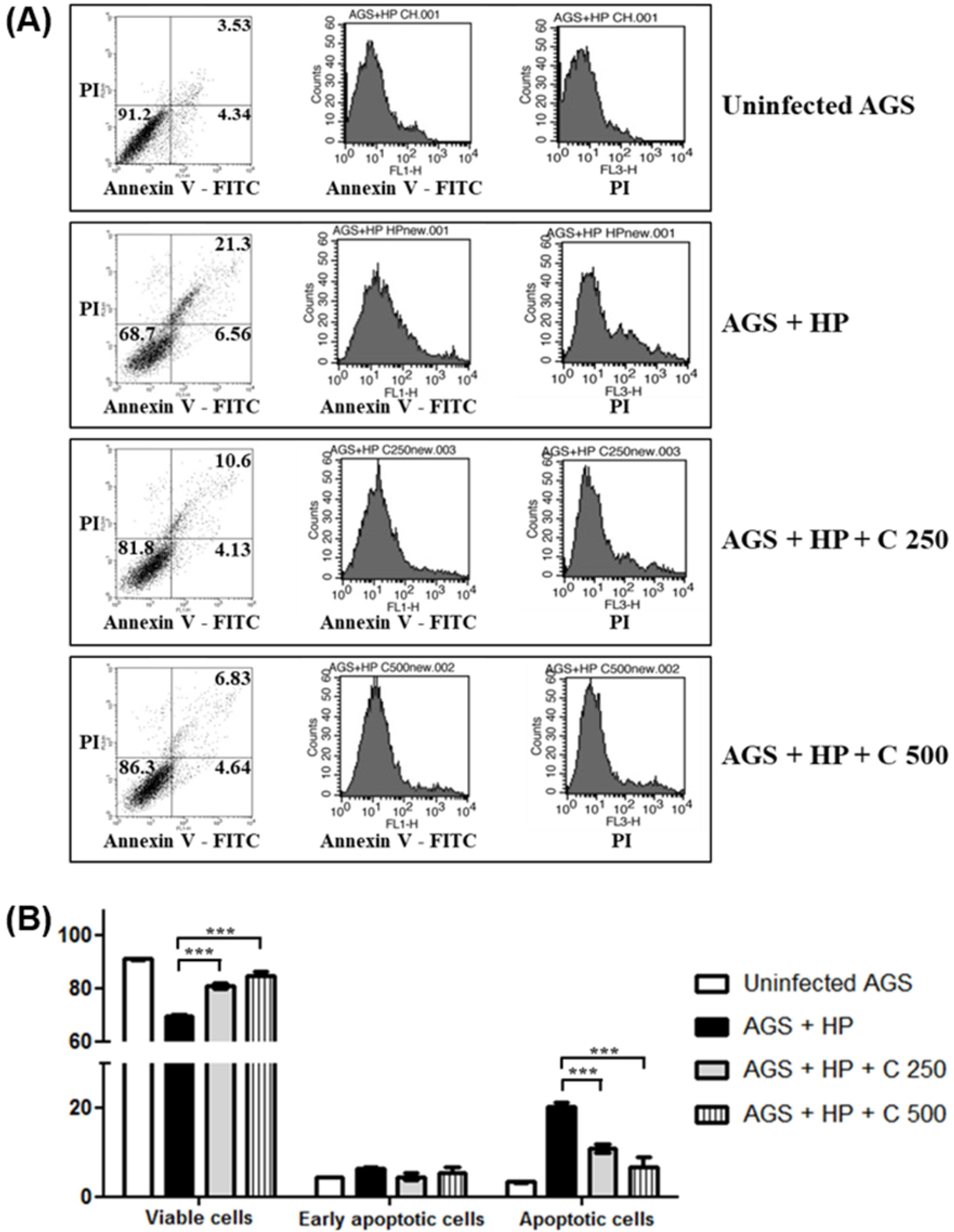
Effect of β-caryophyllene on apoptosis of AGS cell infected with *H. pylori*. AGS cells were infected with *H. pylori* (200 MOI) and treated with indicated dose of β-caryophyllene (250, 500 μg/mL) for 24 h. After incubation, the cells were stained with annexin V-FITC and PI, and subjected to flow cytometry. β-caryophyllene alleviated apoptosis of AGS cells induced by *H. pylori* infection. (A) Stained cells were analyzed and illustrated on the quadrant by CellQuestPro software. (B) Percentage of cells in apoptosis was analyzed and illustrated as a graph. Results from triplicate experiments were analyzed by Student’s *t*-test (\*\*\**P* < 0.001).

### 3. Inhibitory effects of β-caryophyllene on the translocation of bacterial CagA and VacA proteins

Cytoskeletal rearrangement and resultant morphological change so-called hummingbird phenotype is a noted outcome appearing after injection of CagA protein into AGS cells and accumulation of cytoplasmic vesicles (vacuolation) is induced by VacA translocation. Furthermore, it was reported that CagA and VacA proteins are closely associated with the *H. pylori*-induced apoptosis in gastric epithelial cells. AGS cells were infected with *H. pylori* and at the same time treated with β-caryophyllene and then incubated for 24 h. As a result, *H. pylori* infection resulted in the morphological changes of AGS cells such as the hummingbird phenotype and vacuolation in the cytosol. However, the morphological changes were alleviated by β-caryophyllene treatment (Fig 4A). It was confirmed that both CagA and VacA protein levels were increased by *H. pylori* infection but the protein levels were decreased in AGS cells after β-caryophyllene treatment (Fig 4B). The experiment results were corresponded with the morphological results.

**Fig 4.**
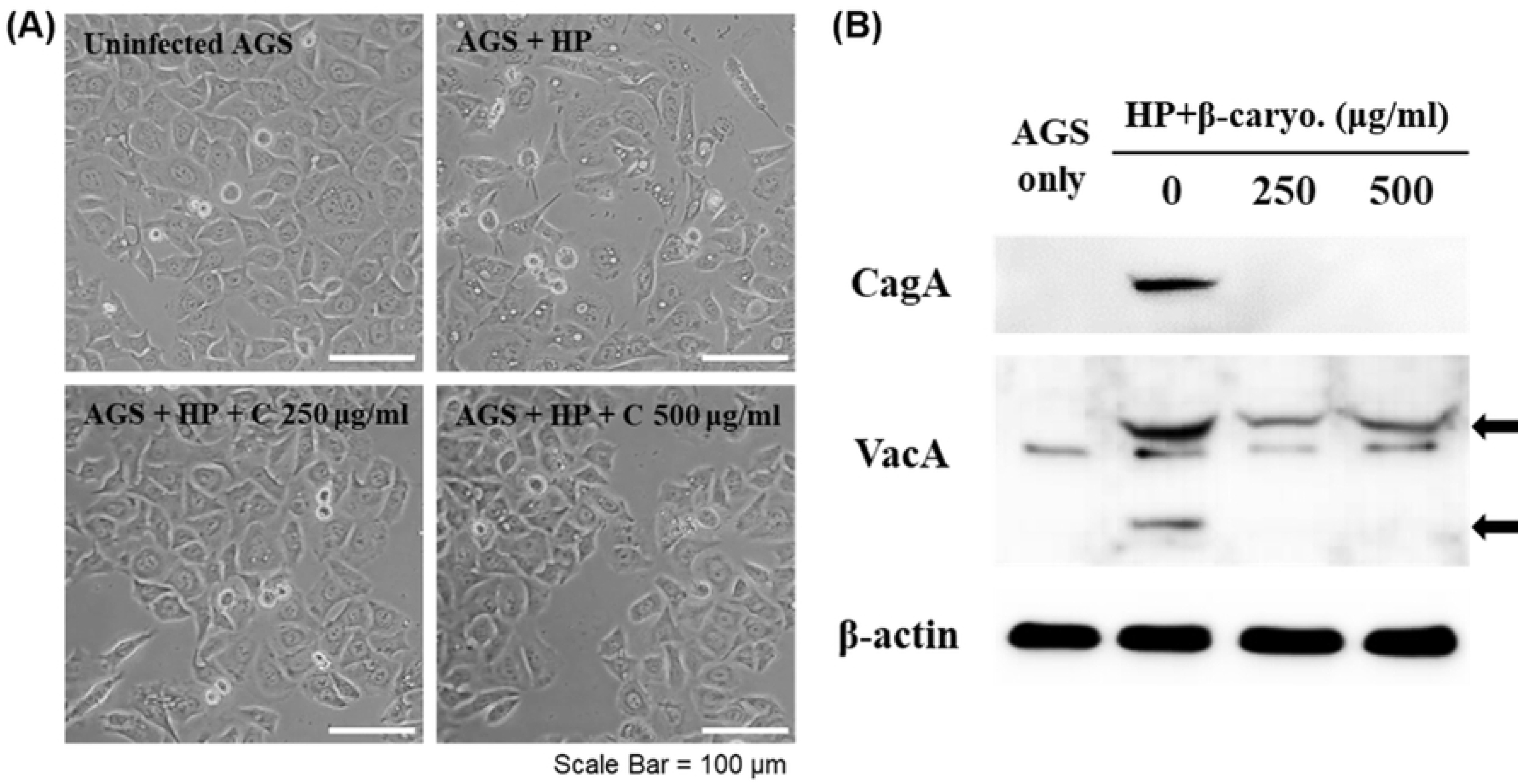
Inhibitory effect of β-caryophyllene on CagA and VacA translocation into AGS cells by *H. pylori*. AGS cells were infected with *H. pylori* (200 MOI) and treated with indicated dose of β-caryophyllene (250, 500 μg/mL) for 24 h. After incubation, (A) morphological changes were observed with an inverted microscope (×200). (B) The cell lysates were assessed by Western blotting to detect translocated CagA and VacA protein to AGS cells.

To elucidate the reasons why the translocation of CagA and VacA proteins to AGS cells was inhibited by caryophllene, we investigated the effect of β-caryophyllene on *H. pylori* directly on the mRNA and protein expressions of CagA and VacA. The mRNA levels of *cag*A and *vac*A genes in *H. pylori* treated with β-caryophyllene were reduced by β-caryophyllene treatment (Fig 5A and 5B). The protein levels of CagA and VacA in *H. pylori*-treated with β-caryophyllene were also downregulated of CagA and VacA in *H. pylori* (Fig 5C and 5D). These results, collectively, suggested that β-caryophyllene inhibited the production of CagA and VacA.

**Fig 5.**
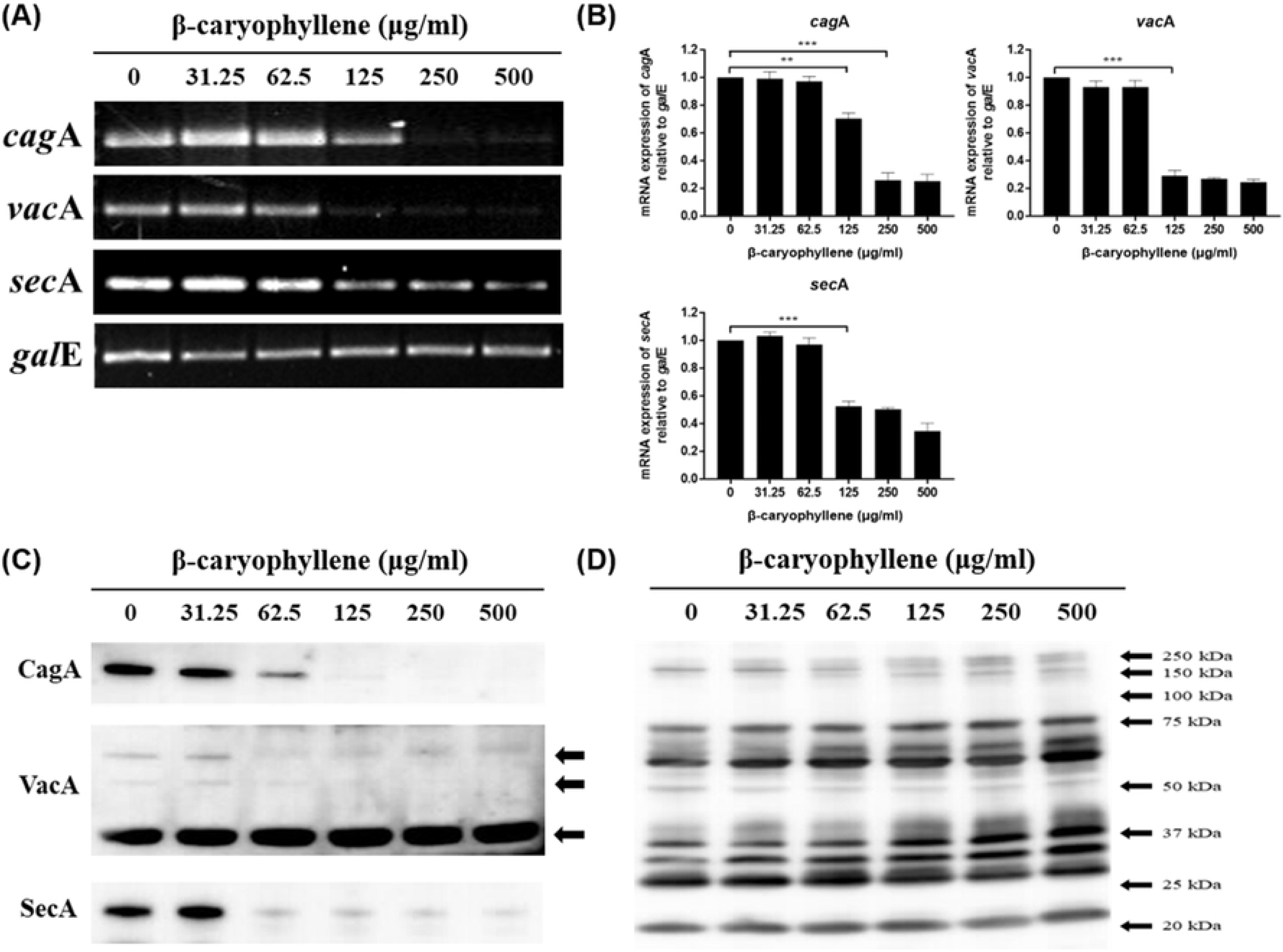
Expression of virulence factors in *H. pylori* treated with β-caryophyllene. *H. pylori* was treated with indicated concentrations of β-caryophyllene in Mueller-Hinton broth for 72 h. (A) The RNA was extracted and subjected to RT-PCR to detect the expression levels of virulence factors. Constitutively expressed gale was used as an internal control. (B) Density of the bands were illustrated as a graph, and the results from triplicate experiments were analyzed by Student’s *t*-test (\*\**P* < 0.01 and \*\*\**P* < 0.001). (C) The bacterial lysates were assessed by Western blotting to detect CagA, VacA and SecA protein. (D) Rabbit anti-*H. pylori* polyclonal antibody was used as an internal control.

Besides, the mRNA and protein levels of SecA in *H. pylori*, an ATPase responsible for the regulation of T5aSS that secrets VacA protein, were suppressed by β-caryophyllene treatment (Fig 5). The mRNA expression levels of *vir*B2, *vir*B4 and *vir*B8, T4SS components in *H. pylori,* were reduced in a β-caryophyllene dose-dependent manner (Fig 6). These results suggest that inhibition of CagA and T4SS expression by β-caryophyllene might be in part associated with the decreased translocation of CagA to AGS cells and downregulated SecA as well as VacA might be induced VacA translocation to AGS cells.

**Fig 6.**
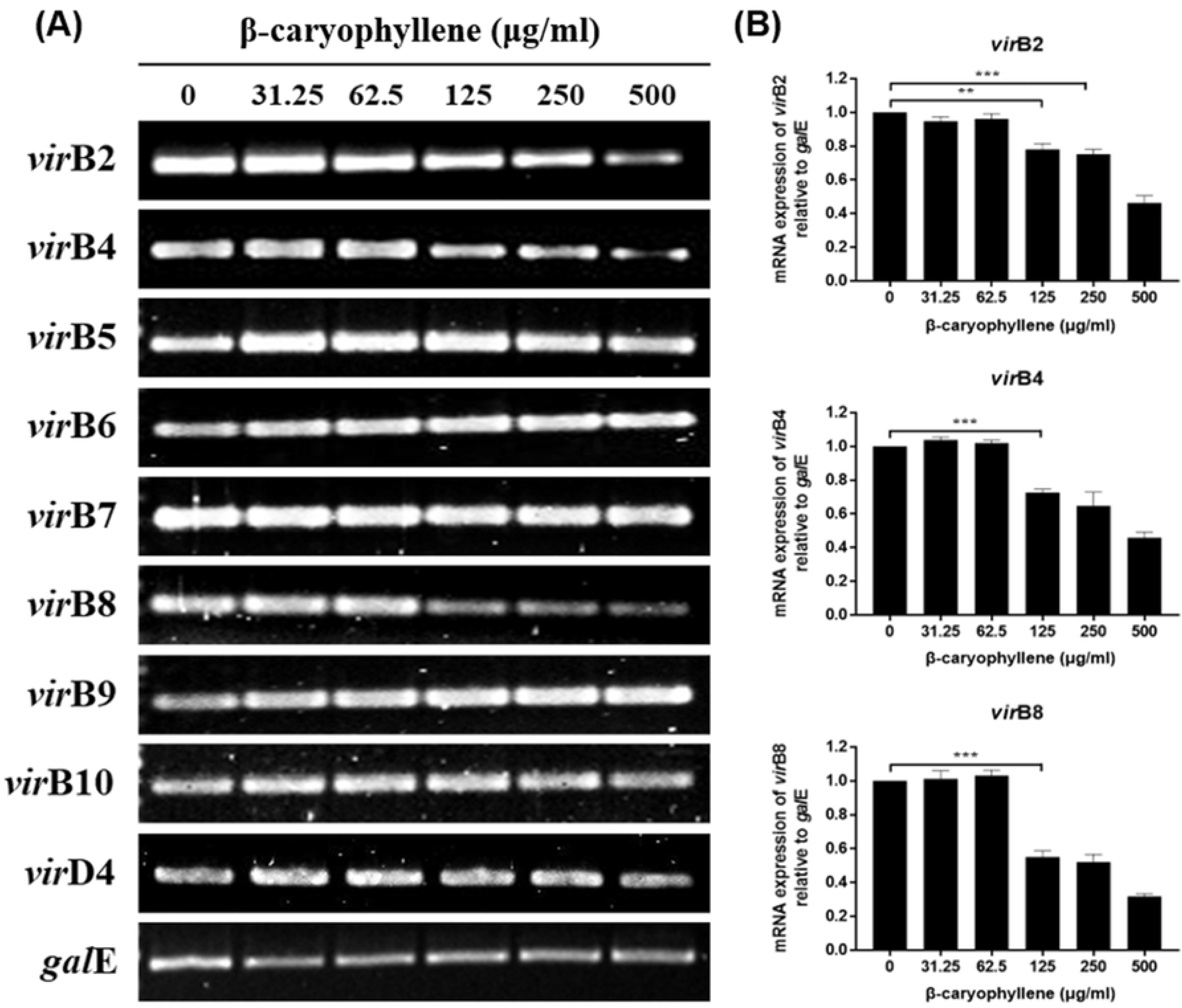
Inhibitory effect of β-caryophyllene on the CagA translocation through type IV secretion system. (A) *H. pylori* was treated with indicated concentrations of β-caryophyllene and the RNA was extracted. The collected RNA was subjected to RT-PCR to detect the expression of T4SS components (*vir*B2, *vir*B4, *vir*B5, *vir*B6, *vir*B7, *vir*B8, *vir*B9 and *vir*D4). The *gal*E was used as an internal control. (B) Density of the bands were illustrated as a graph, and the results from triplicate experiments were analyzed by Student’s *t*-test (\*\**P* < 0.01 and \*\*\**P* < 0.001).

### 4. Therapeutic effects of β-caryophyllene on Mongolian gerbils infected with *H. pylori*

Mongolian gerbils were infected with *H. pylori* to establish an *H. pylori*-infected animal model. The animal model was treated with or without β-caryophyllene to evaluate the therapeutic effects *in vivo*. Animals were divided into four groups. “Normal control group” comprised of 7 gerbils that were inoculated with corn oil. “*H. pylori* control group” comprised of 8 gerbils that were inoculated with *H. pylori* (1 × 10^9^ cells) and were given no further treatment. Group, “High dose” and group, “Low dose” were inoculated with *H. pylori* (1 × 10^9^ cells) and the administration of β-caryophyllene was started at 2 weeks after initial inoculation. β-caryophyllene was diluted with sterile corn oil at two concentrations. Five hundred μg/g was prepared for high dose group, and one hundred μg/g was prepared for low dose group (S1 Appendix).

Mongolian gerbils in each group were sacrificed at 0 week, 6 weeks, and 12 weeks after beginning of the β-caryophyllene treatment. RNA was extracted from stomach of the Mongolian gerbils, and then RT-PCR targeting *H. pylori*’s 16S rRNA was performed to evaluate the presence of living *H. pylori*. The result at 0 week showed successful colonization of *H. pylori* in all the infected groups. Presence of *H. pylori* was detected in all the *H. pylori* infected Mongolian gerbils without β-caryophyllene treatment, and the infection lasted until 12 weeks. However, *H. pylori* was not detected in the groups treated with β-caryophyllene since 6 weeks indicating successful eradication of *H. pylori* by β-caryophyllene (S2 Appendix).

Furthermore, gastric tissue sections of each group were prepared and presence of *H. pylori* was also investigated by using immunohistochemistry (IHC) stain. Anti-*H. pylori* antibody was used to detect the presence of *H. pylori*. There was no positive signal observed in the uninfected gerbil group (Normal control group) (Fig 7 and 8). In contrast, positive signals for anti-*H. pylori* antibody were observed abundantly in gastric mucosal and submucosal layer of β-caryophyllene-untreated group (*H. pylori* control group), which indicated a marked *H. pylori* infection in gastric epithelium of the gerbils (Fig 7 and 8). *H. pylori* were also detected in the β-caryophyllene-treated groups at 0 week, but the positive signals were decreased over times after β-caryophyllene treatment suggesting that both low dose (100 μg/g) and high dose (500 μg/g) of β-caryophyllene significantly reduced the degree of infection by *H. pylori* (Fig 7 and 8). These results collectively indicate the therapeutic effects of β-caryophyllene on Mongolian gerbils infected with *H. pylori*.

**Fig 7.**
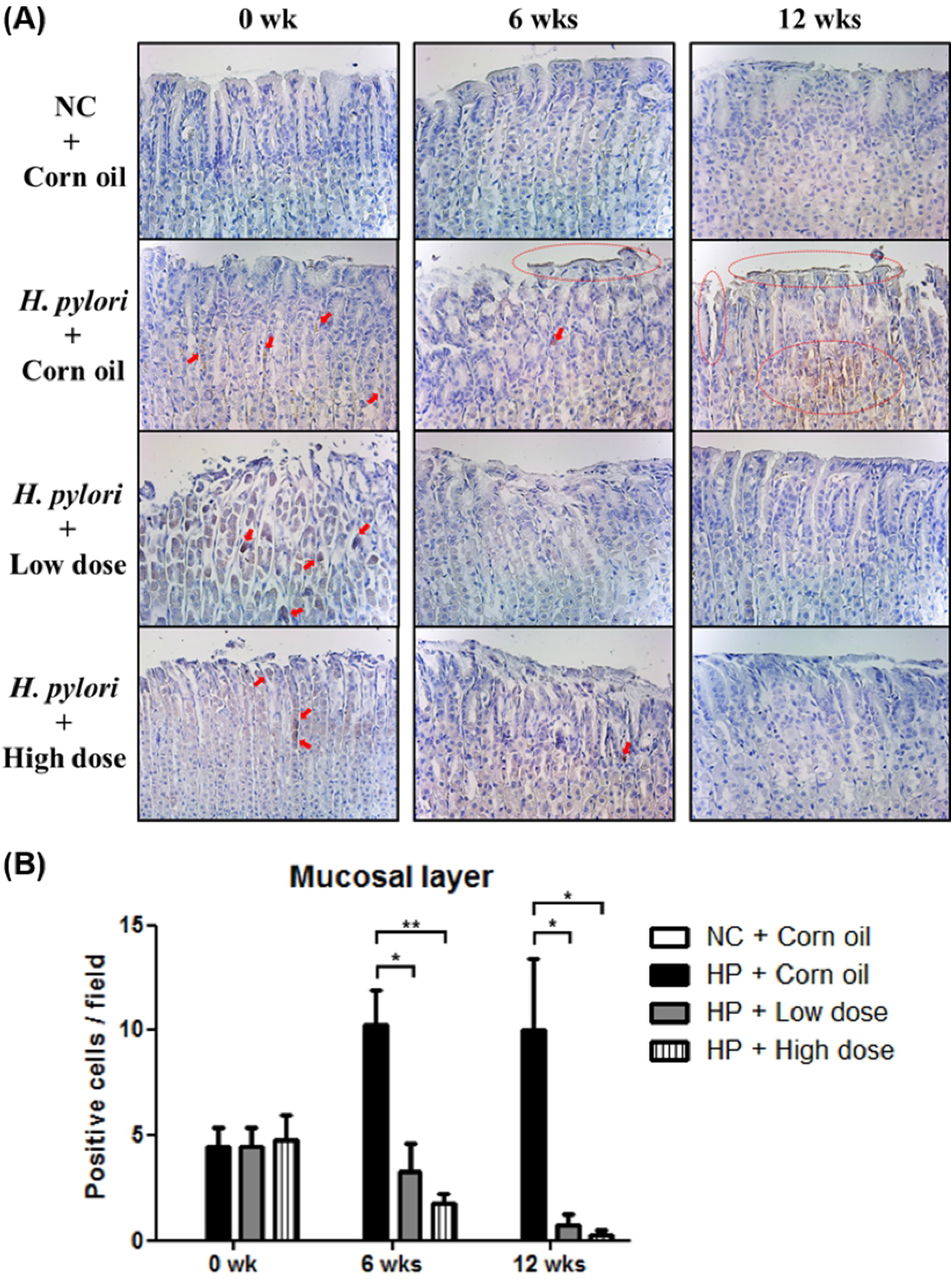
Immunohistochemistry of gastric mucosal layer in *H. pylori*-infected Mongolian gerbils. (A) Photomicrograph shows gastric mucosal layer of *H. pylori*-infected Mongolian gerbils (Magnification ×200). (B) Statistical analysis of data obtained from photomicrograph images. *H. pylori*-positive cells were identified by counting the number of cells staining intensely in five fields from each sample and 2-3 gastric tissues were assessed for each group. Data in the bar graphs are presented as the mean ± standard error of mean. Results were analyzed by Student’s *t*-test (\*\**P* < 0.01 and \*\*\**P* < 0.001).

**Fig 8.**
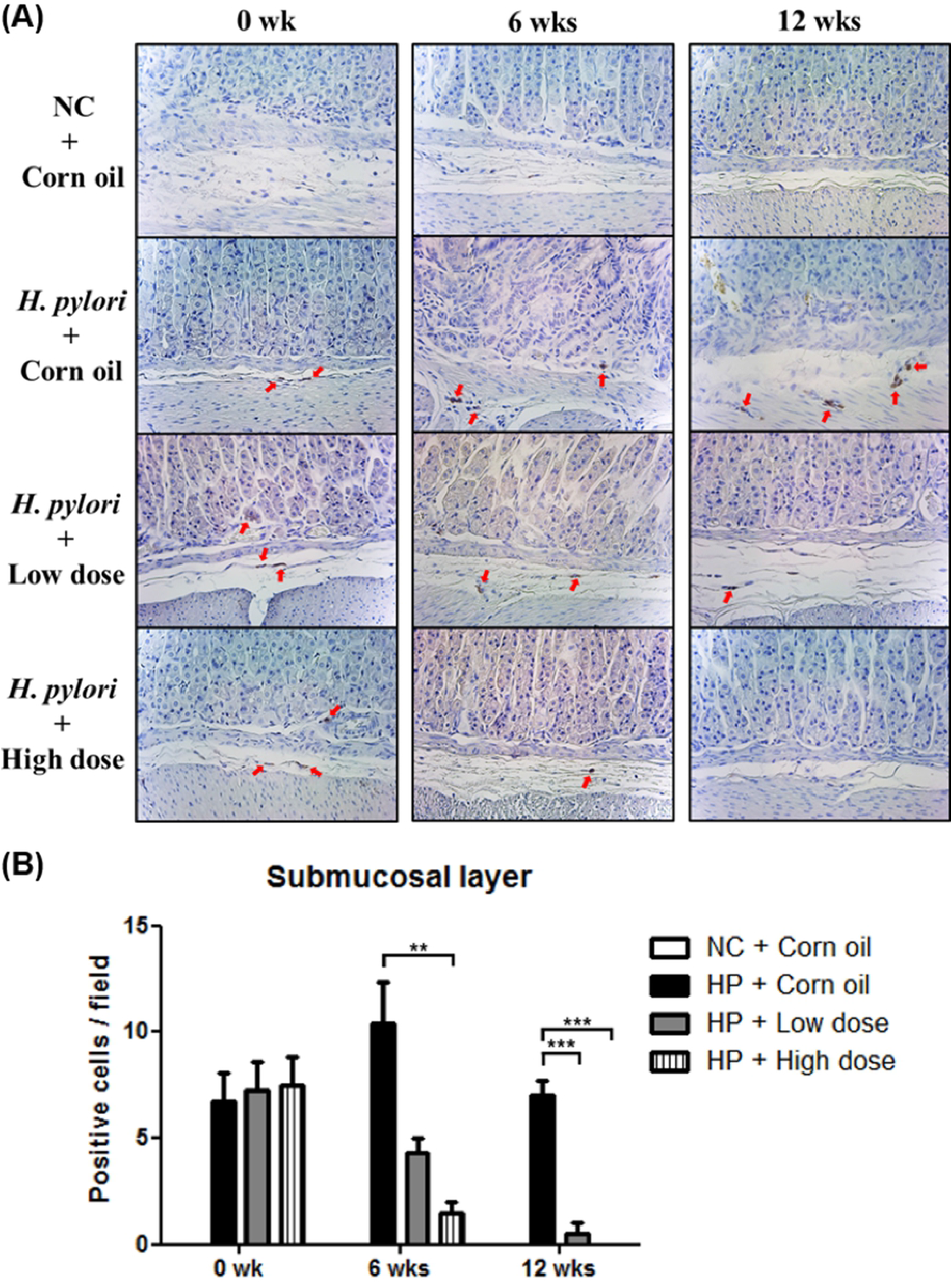
Immunohistochemistry of gastric submucosal layer in *H. pylori*-infected Mongolian gerbils. (A) Photomicrograph shows gastric submucosal layer of *H. pylori*-infected Mongolian gerbils (Magnification ×200). (B) Statistical analysis of data obtained from photomicrograph images. *H. pylori*-positive cells were identified by counting the number of cells staining intensely in five fields from each sample and 2-3 gastric tissues were assessed for each group. Data in the bar graphs are presented as the mean ± standard error of mean. Results were analyzed by Student’s *t*-test (\*\**P* < 0.01 and \*\*\**P* < 0.001).

### 5. Inhibitory effects of β-caryophyllene on the *H. pylori*-induced inflammation

The gastric tissue sections were also stained with hematoxylin & eosin (H&E) and observed with microscope to evaluate the inflammation in the tissue. In the *H. pylori* control group, abnormally disrupted shape of gastric epithelium was observed at 12 weeks indicating that inflammation was occurred by *H. pylori* infection. However, the gastric epithelium was almost undamaged in β-caryophyllene-treated groups suggesting that the inflammation induced by *H. pylori* was alleviated by β-caryophyllene treatment (Fig 9A).

**Fig 9.**
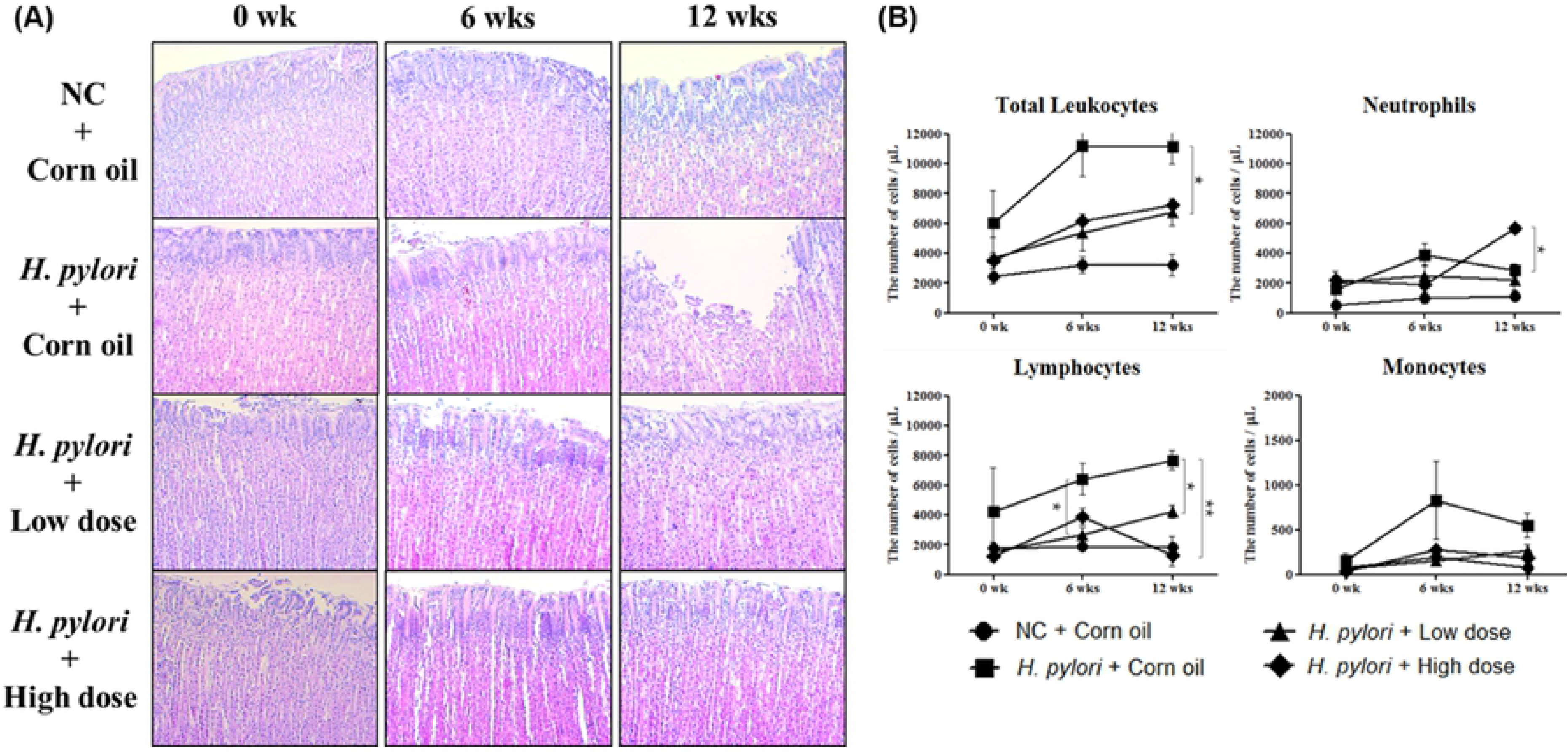
H&E staining of gastric mucosal layer and blood leukocyte count in *H. pylori*-infected Mongolian gerbils. (A) The stomach tissues from *H. pylori*-infected Mongolian gerbils were collected for H&E stain. Photomicrograph shows gastric mucosal layer of *H. pylori*-infected Mongolian gerbils (Magnification ×200). (B) Blood was taken from *H. pylori*-infected Mongolian gerbils at the specified time after β-caryophyllene administration (0, 6, 12 weeks) and subjected to blood cell counting. Data in the bar graphs are presented as mean ± standard error of mean. The results were significant (\**P* < 0.05 and \*\**P* < 0.01) as compared with the *H. pylori* control. Circle, NC + corn oil group; square, *H. pylori* + corn oil group; triangle, *H. pylori* + low dose group; diamond, *H. pylori* + high dose group.

Blood was collected from the left atrium of all the gerbils during sacrifice. In the *H. pylori* control group, number of total leukocytes was dramatically increased since 6 weeks in comparison to the normal control group indicating leukocytosis. Although, slight increases of total leukocytes were also observed in low dose (100 μg/g) or high dose (500 μg/g) of β-caryophyllene treated groups, β-caryophyllene treatment alleviated leukocytosis induced by *H. pylori* infection. Similar pattern was also observed in the number of lymphocytes and monocytes. Lymphocytes and monocytes were also increased in the blood of gerbils by *H. pylori* infection but the increases of lymphocytes and monocytes were alleviated by treatment of β-caryophyllene. However, no significant increase of neutrophils was observed after *H. pylori* infection (Fig 9B).

### 6. Cytotoxicity of β-caryophyllene *in vivo*

Chemicals may have toxicity thus can damage the organs such as liver and kidneys during the medication. Weight of the Mongolian gerbils was measured periodically during the experiment. However, no significant differences were observed among the groups (S1 Table). Hepatotoxicity and nephrotoxicity was evaluated in the gerbils exposed to β-caryophyllene. The hepatic and renal tissue sections of the gerbils were stained with hematoxylin & eosin and observed by microscope. In the results, histopathological lesions were not detected in the low dose (100 μg/g) of β-caryophyllene-treated group. In the high dose (500 μg/g) of β-caryophyllene-treated group, however, hemorrhagic lesions containing red blood cells were observed in both hepatic and renal tissues (Fig 10). These results suggest that low dose (100 μg/g) of β-caryophyllene has no liver toxicity and nephrotoxicity.

**Fig 10.**
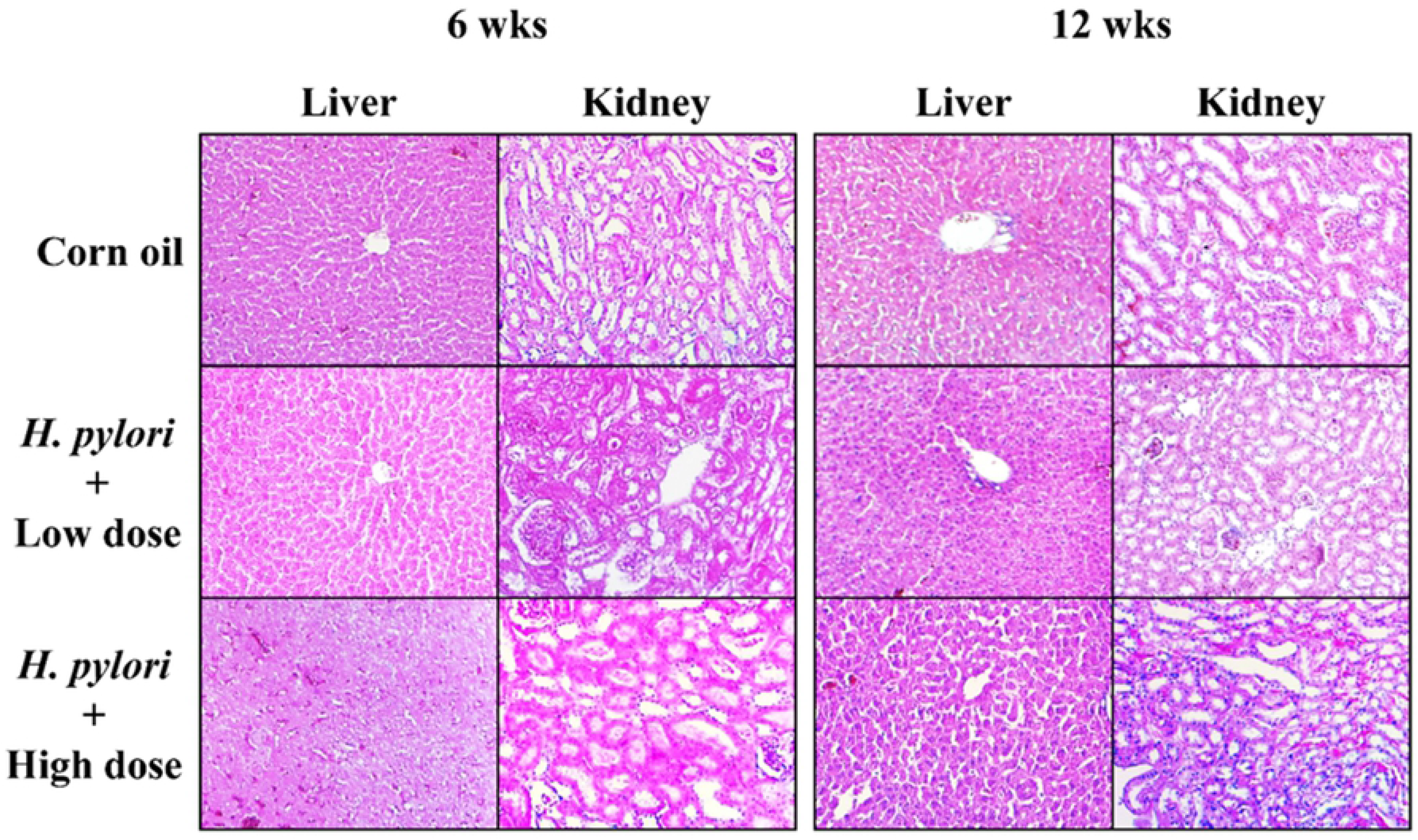
Assessment of toxicity in liver and kidney. Mongolian gerbils were orogastrically administrated β-caryophyllene or corn oil every day for 6 weeks or 12 weeks. A total of 10 Mongolian gerbils were sacrificed at 6 and 12 weeks and their renal and hepatic tissues were excised for histopathologic analysis. Low dose (100 μg/g) of β-caryophyllene has no hepatotoxicity and nephrotoxicity. However, high dose (500 μg/g) of β-caryophyllene has mild hepatotoxicity and nephrotoxicity.

However, high dose (500 μg/g) of β-caryophyllene may damage the organs, thus cautious selection for dose of β-caryophyllene seems to be necessary to avoid toxicity.

## IV. DISCUSSION

Development of new antimicrobial agents with fewer disadvantages is necessary for eradication of *H. pylori*. Β-caryophyllene is one of the natural compounds which easily found in the essential oils of many plants [34–36]. According to several reports, β-caryophyllene has antibacterial effect on cariogenic bacteria and food-spoilage bacteria [39–41]. This study demonstrated the inhibitory effect of β-caryophyllene on *H. pylori* growth and the protective effect against *H. pylori* infection as well as antibacterial therapeutic effect. The aim of this study was to discover the new antimicrobial agent from nature which may inhibit the *H. pylori* infection, thus this study aimed to investigate on the effect of β-caryophyllene as one of the natural compound which possess multiple pharmacological properties [33].

In this study, inhibitory effect of β-caryophyllene on *H. pylori* was firstly evaluated by disc diffusion assay (Fig 1A). Then, it was observed that the MIC of β-caryophyllene against *H. pylori* was 1,000 μg/mL in the broth dilution test (Fig 1B). Expressions of replication genes of *H. pylori* were evaluated to elucidate how β-caryophyllene inhibits the growth of *H. pylori*. β-caryophyllene treatment decreased the mRNA expression levels of *dna*E, *dna*N, *hol*B and *gyr*A genes (Fig 1C and 1D). DnaE is the catalytic α subunit of DNA polymerase III. It has been reported that the mutant *dna*N was unable to support *E. coli* growth [43]. Song *et al.* have shown that *E. coli* strains bearing chromosomal knockout of *hol*B gene was not viable [44]. These studies demonstrated that the DnaE, DnaN and HolB are necessary for cell growth, thus they are indispensable. DNA gyrase is pivotal for the process of bacterial replication, so that it has received the most attention for developing antibiotics such as novobiocin (the ATP site inhibitor) and flioroquinones (the catalytic site inhibitor) [45, 46]. Therefore, interruption of bacterial replication via down-regulation of *dna*E, *dna*N, *hol*B and *gyr*A genes by β-caryophyllene may explain the inhibitory mechanism of β-caryophyllene against *H. pylori* growth.

In this study, it was evaluated whether β-caryophyllene can alleviate the deleterious effects of *H. pylori* infection on gastric epithelial cells. β-caryophyllene showed no cytotoxic effect on AGS cells upto 500 μg/mL (Fig 2A). Since the apoptotic cell death is closely related to the gastric cancer development, the effect of β-caryophyllene treatment on *H. pylori*-induced apoptotic cell death was assessed by performing cell viability test, Western blotting and Annexin V staining [47, 48]. The results revealed that apoptosis induced by *H. pylori* infection was alleviated in AGS cells by β-caryophyllene treatment (Fig 2B, 2C, 2D, 3A and 3B).

Furthermore, β-caryophyllene treatment diminished the hummingbird phenotype and vacuolation on *H. pylori*-infected AGS cells dose dependently (Fig 4A) and also inhibited CagA and VacA translocation into the *H. pylori*-infected AGS cells (Fig 4B). It was due to the reduction of the CagA, VacA and SecA as well as that of several T4SS components including *vir*B2, *vir*B4 and *vir*B8 expression by β-caryophyllene treatment (Fig 5 and 6). Both CagA and VacA proteins disrupt intracellular signaling in host cells that lead to uncontrolled growth of the cells and inflammatory responses [1, 2, 49, 50]. Therefore, the results imply that the decrease of CagA and VacA by β-caryophyllene treatment will alleviate the tumorigenesis and inflammation induced by *H. pylori* in gastric mucosa. Collectively, these *in vitro* data provides evidence supporting that β-caryophyllene may helpful to attenuate the deleterious effects such as hummingbird phenotype, vacuolation and apoptotic cell death induced by *H. pylori* infection.

The *in vivo* experiments in this study demonstrated the therapeutic effects of β-caryophyllene. IHC staining data indicated that β-caryophyllene administration relieves the infection of *H. pylori* (Fig 7 and 8) and H&E staining data showed that β-caryophyllene treatment diminishes inflammation in *H. pylori*-infected stomach tissues (Fig 9). The reason why inflammatory signs are decreased in β-caryophyllene-treated groups is owing to the colonized *H. pylori* might be eradicated by β-caryophyllene administration. In long-term *in vivo* study, Wiedemann *et al*. reported that early inflammation was observed at 8 weeks and precancerous gastric changes were developed at late time point (32 or 64 weeks) [51]. According to these findings, it might be able to detect more severe inflammatory signs and precancerous changes from *H. pylori* control group, if the infection was maintained over 32 or 64 weeks.

In the hematological study, the numbers of total leukocytes in *H. pylori* control group were vertically increased at 6 weeks and maintained highly at 12 weeks. The numbers of total leukocytes in β-caryophyllene-treated groups, however, were gradually increased after β-caryophyllene administration (Fig 9B). Detailed comparison of hematological changes between β-caryophyllene-untreated group and β-caryophyllene-treated groups also showed that the treatment of β-caryophyllene alleviates *H. pylori* infection. Because neutrophils actively resist to bacterial infection, increased-neutrophil count is commonly found in bacterial infection. However, elevated numbers of neutrophils were relieved by β-caryophyllene administration. This finding implies that bacterial infection is dwindled or bacterial pathogenesis is hindered. In addition to neutrophils, the increased number of lymphocytes and monocytes were lessened at 6, 12 weeks after β-caryophyllene administration. Contrary to human leukocytes, the composition ratio of lymphocytes in total leukocytes is over 60% in Mongolian gerbils [52, 53]. Resulting from the ratio, total changes of leukocytes may be mostly affected by the changes of lymphocytes in Mongolian gerbils. These data correspond with preceded histological data and this certainly supports the therapeutic effect of β-caryophyllene on *H. pylori* infection. Meanwhile, there may be controversial about neutrophil count data because of the sudden fluctuation in high dose group at 12 weeks. It is speculated that daily medication with high dose of β-caryophyllene may lead the chemical toxicity.

*In vivo* toxicological studies of β-caryophyllene have been revealed that it has a very large therapeutic window, with an oral LD_50_ of more than 5,000 μg/g in rats and mice [54, 55]. On the other hand, the toxicological effects of β-caryophyllene on Mongolian gerbil are not reported. In this regard, the two doses (100, 500 μg/g) of β-caryophyllene which are less than 1/10 of LD_50_ were chosen to assess the toxicological effects. In this study, mortality or other signs of toxicity were not detected in the low dose (100 μg/g) of β-caryophyllene-treated group. However, mild hepatotoxicity and nephrotoxicity were detected in the high dose (500 μg/g) group. This result conformed with neutrophil count which indicates the damage of organs.

Further studies are required to fully elucidate about the anti-inflammatory and anti-apoptotic mechanism of β-caryophyllene against *H. pylori*. Previous study from Tambe *et al.* suggested that β-caryophyllene has gastric cytoprotective effect [56]. Thus it would be intriguing to study about the anti-inflammatory and anti-apoptotic mechanism of β-caryophyllene during *H. pylori* infection. Along with these mechanism study, the oral LD_50_ of β-caryophyllene in Mongolian gerbil seems to be necessary to establish.

## Materials & Methods

### 1. Bacterial culture and determination of antibacterial activity

*H. pylori* ATCC 49503 strain (East-asian type: CagA^+^/VacA^+^, American Type Culture Collection, Manassas, VA, USA) was grown on Brucella agar plates (Becton-Dickinson, Braintree, MA, USA) supplemented with 10% bovine serum (BRL Life Technologies, Grand Island, NY, USA) at 37°C for 72 h under humidified atmosphere with 5% CO_2_. For disc diffusion method, the number of bacteria in the *H. pylori* suspension was adjusted to McFarland scale 2 (6 × 10^8^ cells/mL) and spread evenly them on Mueller-Hinton agar (Becton-Dickinson) supplemented with 10% bovine serum. β-caryophyllene was provided by SFC BIO Co., Ltd in republic of Korea. The discs which had been impregnated with a series of β-caryophyllene were placed on the plate and then the plate was incubated for 72 hr. The inhibition zone was measured each diameter. To determine the minimum inhibitory concentration (MIC) of β-caryophyllene against *H. pylori*, the number of bacterial particles in the *H. pylori* suspension was set to McFarland scale 0.5 (1.5 × 10^8^ cells/mL). Various concentrations of β-caryophyllene (7.81-4000 μg/mL) were treated and the bacteria were incubated for 72 h and final optical density (600 nm) of the bacterial suspension was measured by using NanoQuant spectrophotometer (infinite M200, TECAN, Männedorf, Switzerland). For normal control, the same volume of ethanol was administrated to culture media.

### 2. Mammalian cell culture

AGS gastric adenocarcinoma cells (ATCC CRL-1739) were cultured in DMEM medium (BRL Life Technologies) supplemented with 10% fetal bovine serum (BRL Life Technologies) and streptomycin-penicillin (100 μg/mL and 100 IU/mL, BRL Life Technologies). Cells were infected with *H. pylori* as a concentration of 200 multiplicity of infection (MOI) without addition of antimicrobial agents in media and then treated with β-caryophyllene (250 and 500 μg/mL). For normal control, the equivalent amount of ethanol was administrated to culture media. In the experimental groups, the results were compared with normal control group and each experiment were repeated over three times to confirm data.

### 3. RT-PCR

*H. pylori* ATCC 49503 strain was grown in Mueller-Hinton broth (Becton-Dickinson) for 72 h. Cultured *H. pylori* was washed twice with phosphate-buffered saline (PBS) and total RNA was extracted using Trizol reagent (Invitrogen, Carlsbad, CA, USA) as described in the manufacturer’s instructions. cDNA was synthesized by reverse transcription with random hexamer (Invitrogen) and Moloney murine leukemia virus reverse-transcriptase (MMLV-RT, Invitrogen). Subsequent PCR amplification performed in a thermocycler using specific primers. The PCR primer sequences used in this study are listed in Table 1 [57–66]. Gel images have been captured and analyzed using the Quantity One system (Bio-Rad, Hercules, CA, USA).

### 4. Western blotting

Bacteria and AGS cells were lysed with radio immunoprecipitation assay (RIPA) lysis buffer (Millipore, Billerica, MA, USA) containing a protease inhibitor cocktail (Calbiochem, San Diego, CA, USA). The cell lysates were incubated on ice for 30 min. In order to lyse the bacterial cells completely, the mixture was sonicated for 2 minutes with 10 second intervals (Sonicator XL-2020, Heat Systems Ultrasonics, Pittsburgh, PA, USA). The cell lysates were then centrifuged at 13,000 *g* for 10 min at 4°C and the supernatants were collected. Protein concentrations were determined based on Lowry method and quantified using NanoQuant spectrophotometer Antibodies to detect CagA, VacA and β-actin were purchased from Santa Cruz Biotechnology (Dallas, TX, USA) and polyclonal antibody against whole *H. pylori* (ATCC 49503) and SecA were produced as previously described [20, 66]. Antibodies to detect PARP was purchased from Cell Signaling Technology (Danvers, MA, USA). Anti-*H. pylori* polyclonal antibody and β–actin were used as an internal control for bacteria and mammalian cell proteins.

### 5. WST cell viability assay using EZ-Cytox

AGS cells (1×10^4^ per well) were plated in 96-well plates. After 24 h, cells were treated with various concentrations of β-caryophyllene. The cells were then incubated for 24 h and subjected to water soluble tetrazolium salt (WST) assay by using EZ-Cytox cell viability assay kit according to manufacturer’s instruction. Ten μL of WST solution was added to the cultured media and incubated for 2 h in the CO_2_ incubator. The absorbance was measured at 450 nm by spectrophotometer.

### 6. Annexin V and PI staining

Annexin V and PI staining was performed by using Annexin V-FITC Apoptosis Detection Kit I (Becton-Dickinson) according to the manufacturer’s instruction. The cultured cells were detached with 0.25% trypsin-EDTA, washed twice with cold PBS, and centrifuged at 3000 rpm for 5 mins. The cells were resuspended in 500 μL of 1X binding buffer at a concentration of 5 × 10^5^ cells/mL and 5 μL of Annexin V-FITC and 5 μL of PI were added to the cell suspension. The mixture was incubated for 10 minutes at 37°C in the dark and analyzed by FACS Caliber flow cytometry (Becton-Dickinson).

### 7. Animal and experimental design

Inbred specific pathogen free (SPF) 5 week-old, male and female Mongolian gerbils for mating were purchased from Central Lab Animals, South Korea. Gerbils used in this study were obtained from 10 breeding cages bred inhouse. Animals at the age of 6-7 week-old (n=31) were gathered and challenged orogastrically three-times over five consecutive days with approximately 1 × 10^9^ cells of viable *H. pylori*. β-caryophyllene was orogastrically administrated every day for 12 weeks. All gerbils were sacrificed using CO_2_ euthanasia at different times post-administration (0, 6, 12 weeks). The stomach was excised, opened along the greater curvature. One half was used for a culture study (reisolation) and extraction of RNA while the other was used for immunohistochemical and histopathological analyses. The blood samples were taken from all sacrificed gerbils for hematological examination.

### 8. Assessment of immunohistochemistry

*H. pylori* were detected in infected gastric tissues by immunohistochemistry using a rabbit anti-*H. pylori* antibody (Abcam, Milton, Cambridge, UK). Positive-staining cells were visualized with diaminobenzidine (DAB) (Vectastain Elite ABC Kit for rabbit; Vector Laboratories, Burlingame, CA, USA) and morphometrically analyzed with Leica DM 2500 microscopy and Leica Application Suite software (version 4.4; Leica microsystems, Heerbrugg, Switzerland). The evaluation of *H. pylori*-positive cells as marker of the infection was performed by counting the *H. pylori*-positive cells distributed in the non-infected control gastric tissue.

### 9. Assessment of histopathology

Paraffin embedded longitudinal sections of antrum and corpus were stained with hematoxylin & eosin (H&E) and evaluated. It was graded for gastritis and mucosal changes and analyzed by a double blind test according to the grading scheme for rodents [51, 67]. To determine toxicity of β-caryophyllene *in vivo*, gerbils were orogastrically administrated β-caryophyllene every day for 6 weeks or 12 weeks. The concentrations of β-caryophyllene were either a “low dose” (100 μg/g) or “high dose” (500 μg/g). A total of 7 Mongolian gerbils were sacrificed at 6 and 12 weeks and their renal and hepatic tissues were excised for histopathologic analysis.

### 10. Statistical analysis

Data in the bar graphs are presented as mean ± standard error of mean (SEM). All the statistical analyses were performed using GraphPad Prism 5.02 software (GraphPad Software, San Diego, CA, USA). All the data were analyzed by unpaired Student’s t-test and *P* < 0.05 was considered to be statistically significant (\**P* < 0.05, \*\**P* < 0.01 and \*\*\**P* < 0.001).

### 11. Ethics statement

All *in vivo* experiments and procedures were approved by the Institutional Animal Care and Use Committee (IACUC) of Yonsei University Wonju Campus (approval number: YWC-150612-1). All works was conducted in compliance with government regulations including Welfare Measures for Animal Protection of Ministry of Agriculture, Food and Rural Affairs in republic of Korea.

## Acknowledgments

None

## Supporting Information captions

**S1 Appendix. Experimental protocol *in vivo*.**

Thirty-one gerbils were divided into four groups. “NC + corn oil” were inoculated corn oil without *H. pylori*. “*H. pylori* + corn oil” inoculated with *H. pylori* (1 × 10^9^ cells) 3 times and were given no further treatment. Group, “*H. pylori* + low dose” and group, “*H. pylori* + high dose” were inoculated with *H. pylori* and β-caryophyllene administration was started 2 weeks after the initial inoculation. The treatment was orogastrically performed during 12-week period; 500 μg/g for high dose group, 100 μg/g for low dose group. All gerbils were sacrificed at specified time of administration (0, 6, 12 weeks).

**S2 Appendix. Detection of *H. pylori* in stomach of Mongolian gerbils using RT-PCR.**

The stomach tissues from *H. pylori* infected Mongolian gerbils were collected for RNA extraction then was subjected to RT-PCR. Colonization of *H. pylori* in stomach of Mongolian gerbils was eradicated after β-caryophyllene administration. *H. pylori*-16S rRNA was used to determine the presence of infection. NC: NC + corn oil; HP: *H. pylori* + corn oil; Low: *H. pylori* + low dose group; High: *H. pylori* + high dose group; PC: Positive control.

**S1 Table.** Changes in body weight of *H. pylori* infected Mongolian gerbils.

| Group | Treatment | Week after inoculation |  |  |  |  |
| --- | --- | --- | --- | --- | --- | --- |
|  |  | 0 | 2 | 4 | 8 | 12 |
| I | NC + Corn oil | 63.4 ± 4.8 | 65.7 ± 3.6 | 68.8 ± 3.3 | 72.0 ± 2.6 | 75.7 ± 1.5 |
| II | *H. pylori* + Corn oil | 54.5 ± 2.7 | 58.6 ± 3.5 | 64.5 ± 4.7 | 68.7 ± 6.3 | 68.7 ± 5.5 |
| III | *H. pylori* + low dose | 58.4 ± 6.7 | 60.5 ± 8.2 | 64.7 ± 9.8 | 69.3 ± 12.0 | 71.5 ± 13.4 |
| IV | *H. pylori* + high dose | 60.5 ± 3.5 | 64.0 ± 3.2 | 67.7 ± 3.4 | 71.4 ± 3.0 | 74.0 ± 2.0 |

## REFERENCES

1. Marshall BJ, Warren JR. Unidentified curved bacilli in the stomach of patients with gastritis and peptic ulceration. Lancet. 1984;1(8390):1311–5.

2. Warren JR, Marshall B. Unidentified curved bacilli on gastric epithelium in active chronic gastritis. Lancet. 1983;1(8336):1273–5.

3. Wang F, Meng W, Wang B, Qiao L. Helicobacter pylori-induced gastric inflammation and gastric cancer. Cancer Lett. 2014;345(2):196–202. https://doi.org/10.1016/j.canlet.2013.08.016

4. Wang CC, Yuan YH, Hunt RH. The association between Helicobacter pylori infection and early gastric cancer: A meta-analysis. Am J Gastroenterol. 2007;102(8):1789–98. https://doi.org/10.1111/j.1572-0241.2007.01335.x

5. Webb PM, Law M, Varghese C, Forman D, Yuan JM, Yu M, et al. Gastric cancer and Helicobacter pylori: a combined analysis of 12 case control studies nested within prospective cohorts. Gut. 2001;49(3):347–53.

6. Huang JQ, Sridhar S, Chen Y, Hunt RH. Meta-analysis of the relationship between Helicobacter pylori seropositivity and gastric cancer. Gastroenterology. 1998;114(6):1169–79. https://doi.org/10.1016/S0016-5085(98)70422-6

7. IARCWorkingGroup. Schistosomes, liver flukes and Helicobacter pylori. IARC Working Group on the Evaluation of Carcinogenic Risks to Humans. Lyon, 7-14 June 1994. IARC Monogr Eval Carcinog Risks Hum. 1994;61:1–241.

8. Tegtmeyer N, Wessler S, Backert S. Role of the cag-pathogenicity island encoded type IV secretion system in Helicobacter pylori pathogenesis. FEBS J. 2011;278(8):1190–202. https://doi.org/10.1111/j.1742-4658.2011.08035.x

9. Kwok T, Zabler D, Urman S, Rohde M, Hartig R, Wessler S, et al. Helicobacter exploits integrin for type IV secretion and kinase activation. Nature. 2007;449(7164):862–6. https://doi.org/10.1038/nature06187

10. Merino E, Flores-Encarnacion M, Aguilar-Gutierrez GR. Functional interaction and structural characteristics of unique components of Helicobacter pylori T4SS. FEBS J. 2017. https://doi.org/10.1111/febs.14092

11. Tammer I, Brandt S, Hartig R, Konig W, Backert S. Activation of Abl by Helicobacter pylori: A novel kinase for CagA and crucial mediator of host cell scattering. Gastroenterology. 2007;132(4):1309–19. https://doi.org/10.1053/j.gastro.2007.01.050

12. Poppe M, Feller SM, Romer G, Wessler S. Phosphorylation of Helicobacter pylori CagA by c-Abl leads to cell motility. Oncogene. 2007;26(24):3462–72. https://doi.org/10.1038/sj.onc.1210139

13. Selbach M, Moese S, Hauck CR, Meyer TF, Backert S. Src is the kinase of the Helicobacter pylori CagA protein in vitro and in vivo. J Biol Chem. 2002;277(9):6775–8. https://doi.org/10.1074/jbc.C100754200

14. Zhang XS, Tegtmeyer N, Traube L, Jindal S, Perez-Perez G, Sticht H, et al. A Specific A/T Polymorphism in Western Tyrosine Phosphorylation B-Motifs Regulates Helicobacter pylori CagA Epithelial Cell Interactions. Plos Pathog. 2015;11(2). https://doi.org/10.1371/journal.ppat.1004621

15. Glowinski F, Holland C, Thiede B, Jungblut PR, Meyer TF. Analysis of T4SS-induced signaling by H. pylori using quantitative phosphoproteomics. Front Microbiol. 2014;5. https://doi.org/10.3389/fmicb.2014.00356

16. El-Etr SH, Mueller A, Tompkins LS, Falkow S, Merrell DS. Phosphorylation-independent effects of CagA during interaction between Helicobacter pylori and T84 polarized monolayers. J Infect Dis. 2004;190(8):1516–23. https://doi.org/10.1086/424526

17. Sgouras DN, Trang TT, Yamaoka Y. Pathogenesis of Helicobacter pylori Infection. Helicobacter. 2015;20 Suppl 1:8–16. https://doi.org/10.1111/hel.12251

18. Boquet P, Ricci V. Intoxication strategy of Helicobacter pylori VacA toxin. Trends Microbiol. 2012;20(4):165–74. https://doi.org/10.1016/j.tim.2012.01.008

19. Leyton DL, Rossiter AE, Henderson IR. From self sufficiency to dependence: mechanisms and factors important for autotransporter biogenesis. Nature reviews Microbiology. 2012;10(3):213–25. https://doi.org/10.1038/nrmicro2733

20. Kim SH, Woo H, Park M, Rhee KJ, Moon C, Lee D, et al. Cyanidin 3-O-glucoside reduces Helicobacter pylori VacA-induced cell death of gastric KATO III cells through inhibition of the SecA pathway. International journal of medical sciences. 2014;11(7):742–7. https://doi.org/10.7150/ijms.7167

21. Memon AA, Hussein NR, Miendje Deyi VY, Burette A, Atherton JC. Vacuolating cytotoxin genotypes are strong markers of gastric cancer and duodenal ulcer-associated Helicobacter pylori strains: a matched case-control study. J Clin Microbiol. 2014;52(8):2984–9. https://doi.org/10.1128/JCM.00551-14

22. Li Y, Wandinger-Ness A, Goldenring JR, Cover TL. Clustering and redistribution of late endocytic compartments in response to Helicobacter pylori vacuolating toxin. Mol Biol Cell. 2004;15(4):1946–59. https://doi.org/10.1091/mbc.E03-08-0618

23. Zheng PY, Jones NL. Helicobacter pylori strains expressing the vacuolating cytotoxin interrupt phagosome maturation in macrophages by recruiting and retaining TACO (coronin 1) protein. Cell Microbiol. 2003;5(1):25–40.

24. Molinari M, Salio M, Galli C, Norais N, Rappuoli R, Lanzavecchia A, et al. Selective inhibition of Ii-dependent antigen presentation by Helicobacter pylori toxin VacA. J Exp Med. 1998;187(1):135–40.

25. Nitharwal RG, Verma V, Dasgupta S, Dhar SK. Helicobacter pylori chromosomal DNA replication: current status and future perspectives. FEBS Lett. 2011;585(1):7–17. https://doi.org/10.1016/j.febslet.2010.11.018

26. Swart JR, Griep MA. Primer synthesis kinetics by Escherichia coli primase on single-stranded DNA templates. Biochemistry. 1995;34(49):16097–106.

27. Rowen L, Kornberg A. Primase, the dnaG protein of Escherichia coli. An enzyme which starts DNA chains. J Biol Chem. 1978;253(3):758–64.

28. Diaconu S, Predescu A, Moldoveanu A, Pop CS, Fierbinteanu-Braticevici C. Helicobacter pylori infection: old and new. J Med Life. 2017;10(2):112–7.

29. Ghotaslou R, Leylabadlo HE, Asl YM. Prevalence of antibiotic resistance in Helicobacter pylori: A recent literature review. World J Methodol. 2015;5(3):164–74. https://doi.org/10.5662/wjm.v5.i3.164

30. Wu IT, Chuah SK, Lee CH, Liang CM, Lu LS, Kuo YH, et al. Five-year sequential changes in secondary antibiotic resistance of Helicobacter pylori in Taiwan. World J Gastroenterol. 2015;21(37):10669–74. https://doi.org/10.3748/wjg.v21.i37.10669

31. Zhang YX, Zhou LY, Song ZQ, Zhang JZ, He LH, Ding Y. Primary antibiotic resistance of Helicobacter pylori strains isolated from patients with dyspeptic symptoms in Beijing: a prospective serial study. World J Gastroenterol. 2015;21(9):2786–92. https://doi.org/10.3748/wjg.v21.i9.2786

32. Kao CY, Sheu BS, Wu JJ. Helicobacter pylori infection: An overview of bacterial virulence factors and pathogenesis. Biomed J. 2016;39(1):14–23. https://doi.org/10.1016/j.bj.2015.06.002

33. Sharma C, Al Kaabi JM, Nurulain SM, Goyal SN, Kamal MA, Ojha S. Polypharmacological Properties and Therapeutic Potential of beta-Β-caryophyllene: A Dietary Phytocannabinoid of Pharmaceutical Promise. Curr Pharm Des. 2016;22(21):3237–64.

34. Kamatou GP, Vermaak I, Viljoen AM. Eugenol--from the remote Maluku Islands to the international market place: a review of a remarkable and versatile molecule. Molecules. 2012;17(6):6953–81. https://doi.org/10.3390/molecules17066953

35. Ghelardini C, Galeotti N, Di Cesare Mannelli L, Mazzanti G, Bartolini A. Local anaesthetic activity of beta-β-caryophyllene. Farmaco. 2001;56(5-7):387–9.

36. de Almeida Borges VR, Ribeiro AF, de Souza Anselmo C, Cabral LM, de Sousa VP. Development of a high performance liquid chromatography method for quantification of isomers beta-β-caryophyllene and alpha-humulene in copaiba oleoresin using the Box-Behnken design. J Chromatogr B Analyt Technol Biomed Life Sci. 2013;940:35–41. https://doi.org/10.1016/j.jchromb.2013.09.024

37. Gertsch J. Anti-inflammatory cannabinoids in diet: Towards a better understanding of CB(2) receptor action? Commun Integr Biol. 2008;1(1):26–8.

38. Gertsch J, Leonti M, Raduner S, Racz I, Chen JZ, Xie XQ, et al. Beta-β-caryophyllene is a dietary cannabinoid. P Natl Acad Sci USA. 2008;105(26):9099–104. https://doi.org/10.1073/pnas.0803601105

39. Crevelin EJ, Caixeta SC, Dias HJ, Groppo M, Cunha WR, Martins CH, et al. Antimicrobial Activity of the Essential Oil of Plectranthus neochilus against Cariogenic Bacteria. Evid Based Complement Alternat Med. 2015;2015:102317. https://doi.org/10.1155/2015/102317

40. Schmidt E, Bail S, Friedl SM, Jirovetz L, Buchbauer G, Wanner J, et al. Antimicrobial activities of single aroma compounds. Nat Prod Commun. 2010;5(9):1365–8.

41. Venturi CR, Danielli LJ, Klein F, Apel MA, Montanha JA, Bordignon SA, et al. Chemical analysis and in vitro antiviral and antifungal activities of essential oils from Glechon spathulata and Glechon marifolia. Pharm Biol. 2015;53(5):682–8. https://doi.org/10.3109/13880209.2014.936944

42. Nikolic M, Stojkovic D, Glamoclija J, Ciric A, Markovic T, Smiljkovic M, et al. Could essential oils of green and black pepper be used as food preservatives? J Food Sci Tech Mys. 2015;52(10):6565–73. https://doi.org/10.1007/s13197-015-1792-5

43. Heltzel JM, Scouten Ponticelli SK, Sanders LH, Duzen JM, Cody V, Pace J, et al. Sliding clamp-DNA interactions are required for viability and contribute to DNA polymerase management in Escherichia coli. J Mol Biol. 2009;387(1):74–91. https://doi.org/10.1016/j.jmb.2009.01.050

44. Song MS, Pham PT, Olson M, Carter JR, Franden MA, Schaaper RM, et al. The delta and delta’ subunits of the DNA polymerase III holoenzyme are essential for initiation complex formation and processive elongation. J Biol Chem. 2001;276(37):35165–75. https://doi.org/10.1074/jbc.M100389200

45. de Souza Mendes C, de Souza Antunes AM. Pipeline of Known Chemical Classes of Antibiotics. Antibiotics (Basel). 2013;2(4):500–34. https://doi.org/10.3390/antibiotics2040500

46. Anderle C, Stieger M, Burrell M, Reinelt S, Maxwell A, Page M, et al. Biological activities of novel gyrase inhibitors of the aminocoumarin class. Antimicrob Agents Chemother. 2008;52(6):1982–90. https://doi.org/10.1128/AAC.01235-07

47. Suerbaum S, Michetti P. Medical progress: Helicobacter pylori infection. New Engl J Med. 2002;347(15):1175–86. https://doi.org/10.1056/NEJMra020542

48. Cover TL, Blaser MJ. Helicobacter pylori in health and disease. Gastroenterology. 2009;136(6):1863–73. https://doi.org/10.1053/j.gastro.2009.01.073

49. de Martel C, Ferlay J, Franceschi S, Vignat J, Bray F, Forman D, et al. Global burden of cancers attributable to infections in 2008: a review and synthetic analysis. Lancet Oncol. 2012;13(6):607–15. https://doi.org/10.1016/S1470-2045(12)70137-7

50. Goh KL, Chan WK, Shiota S, Yamaoka Y. Epidemiology of Helicobacter pylori infection and public health implications. Helicobacter. 2011;16 Suppl 1:1–9. https://doi.org/10.1111/j.1523-5378.2011.00874.x

51. Wiedemann T, Loell E, Mueller S, Stoeckelhuber M, Stolte M, Haas R, et al. Helicobacter pylori cag-Pathogenicity island-dependent early immunological response triggers later precancerous gastric changes in Mongolian gerbils. PLoS One. 2009;4(3):e4754. https://doi.org/10.1371/journal.pone.0004754

52. Dillon WG, Glomski CA. The Mongolian gerbil: qualitative and quantitative aspects of the cellular blood picture. Lab Anim. 1975;9(4):283–7. https://doi.org/10.1258/002367775780957250

53. Rosa RA, Glomski CA. The Mongolian gerbil (Meriones unguiculatus): its histological and haematological response to methylcellulose. Lab Anim. 1981;15(2):131–6. https://doi.org/10.1258/002367781780958865

54. Molina-Jasso D, Alvarez-Gonzalez I, Madrigal-Bujaidar E. Clastogenicity of beta-β-caryophyllene in mouse. Biol Pharm Bull. 2009;32(3):520–2.

55. Hart E, Wong L. Acute oral toxicity studies in rats, acute dermal toxicity and primary skin irritation studies in rabbits of 17 fragrance materials. Bionetics Research Laboratories July. 1971;30:1971.

56. Tambe Y, Tsujiuchi H, Honda G, Ikeshiro Y, Tanaka S. Gastric cytoprotection of the non-steroidal anti-inflammatory sesquiterpene, beta-β-caryophyllene. Planta Med. 1996;62(5):469–70. https://doi.org/10.1055/s-2006-957942

57. Kwon DH, Osato MS, Graham DY, El-Zaatari FA. Quantitative RT-PCR analysis of multiple genes encoding putative metronidazole nitroreductases from Helicobacter pylori. Int J Antimicrob Agents. 2000;15(1):31–6.

58. Lee MH WH, Park M, Moon C, Eom YB, Kim SH, Kim JB. Plumbagin Inhibits Expression of Virulence Factors and Growth of Helicobacter pylori. Microbiology and Biotechnology Letters. 2016;44(2):218–26.

59. Wang J, Wang WH, Li J, Liu FX. Celecoxib inhibits Helicobacter pylori colonization-related factors. World J Gastroenterol. 2010;16(7):846–53.

60. Tharmalingam N, Kim SH, Park M, Woo HJ, Kim HW, Yang JY, et al. Inhibitory effect of piperine on Helicobacter pylori growth and adhesion to gastric adenocarcinoma cells. Infect Agent Cancer. 2014;9(1):43. https://doi.org/10.1186/1750-9378-9-43

61. Clayton C, Kleanthous K, Tabaqchali S. Detection and Identification of Helicobacter-Pylori by the Polymerase Chain-Reaction. J Clin Pathol. 1991;44(6):515–6. https://doi.org/DOI10.1136/jcp.44.6.515

62. Boonjakuakul JK, Canfield DR, Solnick JV. Comparison of Helicobacter pylori virulence gene expression in vitro and in the rhesus macaque. Infect Immun. 2005;73(8):4895–904. https://doi.org/10.1128/Iai.73.8.4895-4904.2005

63. Niehues M, Stark T, Keller D, Hofmann T, Hensel A. Antiadhesion as a functional concept for prevention of pathogens: N-Phenylpropenoyl-L-amino acid amides as inhibitors of the Helicobacter pylori BabA outer membrane protein. Mol Nutr Food Res. 2011;55(7):1104–17. https://doi.org/10.1002/mnfr.201000548

64. Kim JB, Kim GH, Kim H, Jin HS, Kim YS, Ha SH, et al. A Novel PCR Primers HPU185 and HPL826 Based on 16S rRNA Gene for Detection of Helicobacter pylori. The Journal of the Korean Society for Microbiology. 2000;35(4):283–8.

65. Petry A, Djordjevic T, Weitnauer M, Kietzmann T, Hess J, Gorlach A. NOX2 and NOX4 mediate proliferative response in endothelial cells. Antioxid Redox Signal. 2006;8(9-10):1473–84. https://doi.org/10.1089/ars.2006.8.1473

66. Kim SH, Park M, Woo H, Tharmalingam N, Lee G, Rhee KJ, et al. Inhibitory effects of anthocyanins on secretion of Helicobacter pylori CagA and VacA toxins. International journal of medical sciences. 2012;9(10):838–42. https://doi.org/10.7150/ijms.5094

67. Garhart CA, Redline RW, Nedrud JG, Czinn SJ. Clearance of Helicobacter pylori Infection and Resolution of Postimmunization Gastritis in a Kinetic Study of Prophylactically Immunized Mice. Infect Immun. 2002;70(7):3529–38.

